# Coordinated adaptive changes in insulin, insulin receptor, and inceptor genes in hystricognath rodents

**DOI:** 10.1101/2025.05.26.656141

**Authors:** Maite Hilario, Melisa E. Magallanes, Juan C. Opazo, Enrique P. Lessa

## Abstract

Insulin is a central regulator of glycemia and is highly conserved across mammals. However, hystricognath rodents represent a notable exception, exhibiting insulin proteins that diverge significantly in both sequence and physiological function. Previous studies have shown that this divergence is partially driven by positive selection acting on residues involved in the second binding site of the hormone and in hexamer formation, which is critical for insulin storage in mammals. Inceptor (encoded by the gene *ELAPOR1*) is a recently discovered regulator of the insulin pathway that interacts with proinsulin, insulin, the Insulin receptor (Insr), and the Insulin-like growth factor 1 receptor (Igf1r). Inceptor functions within pancreatic β-cells, where it promotes clathrin-mediated endocytosis of Insr and directs cytoplasmic insulin and proinsulin to lysosomal degradation. Using a comprehensive dataset of mammalian sequences, we tested for positive selection in the genes *Ins*, *Insr*, *Igf1r*, and *ELAPOR1* using maximum likelihood models. Significant signals of positive selection were detected in hystricognath rodents for *Ins*, *Insr*, and *ELAPOR1*, but not for *Igf1r*. In *Ins*, positively selected sites were concentrated in the second binding site and in regions involved in hexamer formation, along with an additional site in the C-peptide. In *Insr*, selected sites were primarily located in the ectodomain, particularly in regions that interact with the second binding site of the insulin molecule. In *ELAPOR1*, selected sites were concentrated in the signal peptide and at the boundary between the second cysteine-rich domain and the mannose 6-phosphate receptor region. Together, these findings provide strong evidence that natural selection has shaped multiple components of the insulin signaling pathway, contributing to the functional divergence observed in hystricognath rodents.

## Introduction

Glycemic regulation in mammals is typically a conserved physiological system involving a network of hormones, growth factors, and receptors. Insulin, secreted by pancreatic β-cells, is a well-characterized hormone responsible for lowering blood glucose levels, and its physiological properties are highly conserved across mammals (Chan and Steiner 2000; Conlon 2001; De Meyts 2015; Rostène and De Meyts 2021). A notable exception to this conservatism is found in hystricognath rodents (infraorder Hystricognathi, (see D’Elía et al. 2019), a diverse group that includes Old World mole-rats (families Bathyergidae and Heterocephalidae), porcupines (Hystricidae), and several New World caviomorph families such as guinea pigs and capybaras (Caviidae), spiny rats (Echimyidae), chinchillas (Chinchillidae), and tuco-tucos (Ctenomyidae).

Given the evolutionary conservation of insulin in mammals (Conlon 2001), the discovery that insulin from guinea pigs (*Cavia porcellus*) and other hystricognaths displays unusual biological properties was unexpected. Early studies showed that guinea pig insulin, and that of other hystricognath species, was not neutralized by anti-bovine insulin antibodies (Davidson and Haist 1965; Davidson et al. 1968; Neville et al. 1973). Further experimental work demonstrated that hystricognath insulins exhibit reduced receptor-binding affinity (Zimmerman et al. 1974; Horuk et al. 1979; Bajaj et al. 1986) and lack the ability to form hexamers, the typical storage form of insulin in the β-cells of most mammals (Wood et al. 1975; Bajaj et al. 1986). These distinct physiological characteristics are at least partially attributable to differences in the primary structure of hystricognath insulin (Bajaj et al. 1986; Opazo et al. 2005). Despite these divergences, hystricognaths can maintain glycemic homeostasis (Opazo et al. 2004), albeit with elevated circulating insulin levels that likely compensate for reduced biological activity. Interestingly, their proinsulins also show enhanced mitogenic activity and interact with additional growth receptors (King and Kahn 1981; King et al. 1983).

Opazo et al. (2005) identified codons in the insulin gene of caviomorph rodents that have evolved under positive selection, likely contributing to their distinct insulin properties. These molecular changes affect the second binding site, essential for insulin-receptor interaction (De Meyts 2015; Lawrence 2021; Choi and Bai 2023), and may also alter regions involved in insulin-insulin interactions, potentially explaining the inability to form hexamers. Adding to the picture of divergence (Irwin 2020), reported accelerated amino acid substitution rates in both the insulin receptor (*Insr*) and the glucagon (*Gcg*) genes in hystricognaths, suggesting broader evolutionary changes across the glycemic regulation axis in this group.

A recent addition to this regulatory network is the protein Inceptor (insulin receptor regulator), encoded by the gene *ELAPOR1* (Endosome-Lysosome Associated Apoptosis and Autophagy Regulator 1). This protein plays a key role in insulin signaling modulation within pancreatic β-cells (Ansarullah et al. 2021). Inceptor interacts with the insulin and Igf1 receptors at the cell membrane to promote clathrin-mediated endocytosis, thereby reducing β-cell responsiveness to insulin. The protein was named “Inceptor” to highlight its role as an insulin receptor regulator. More recently, Siehler et al. (2024) showed that Inceptor also acts as a sorting receptor in the cytoplasm, directing insulin and proinsulin toward lysosomal degradation. While much remains to be learned about this protein, its discovery opens new avenues for understanding insulin regulation in pancreatic β cells (Burchfield and James 2021; Hall and Choi 2021; Kulkarni 2021; Starling 2021) and potentially in other tissues (Bilekova et al. 2023). Given the unique insulin biology of hystricognath rodents, it is plausible that Inceptor may also have undergone lineage-specific modifications that warrant further investigation.

Building on documented interactions among insulin, Insulin receptor, Igf1 receptor, and Inceptor, we hypothesized that the corresponding genes have evolved under positive selection in hystricognath rodents. To test this, we manually annotated these four genes in high-quality genomes from hystricognath species, along with selected rodent and placental mammal genomes, and supplemented them with previously annotated sequence data. Our results confirm the action of positive selection on *Ins* and reveal that *Insr* and *ELAPOR1* have also undergone adaptive evolution in hystricognaths, whereas *Igf1r* has not. Specifically, these genes exhibit elevated nonsynonymous to synonymous substitution ratios (*d_N_/d_S_*) compared to other mammals, and we identify codon sites likely shaped by directional selection. Notably, evolutionary changes concentrate in the second insulin binding site and the corresponding binding regions of *Insr*, highlighting these domains as targets of adaptive molecular evolution in hystricognath rodents.

## Results

### Tests of positive selection

The taxa and tree used for analysis of positive selection are presented in Fig. 1. Replicate runs of both branch and site models, starting with different values of ω, consistently converged on highly similar results. For instance, log-likelihood (lnL) values across replicates were similar to the fifth decimal place. For *Ins*, *Insr*, and *ELAPOR1*, branch models estimating two distinct ω values—one for hystricognath rodents and one for all other mammals—fit the data significantly better than models assuming a single ω across all taxa (*p* < 0.00001 in all cases). In contrast, *Igf1r* showed no significant differences when allowing separate ω estimates for hystricognaths versus other mammals (*p* > 0.97; Table 1). Notably, for *Ins*, *Insr*, and *ELAPOR1*, the ω values were consistently higher in hystricognath rodents, indicating an accelerated rate of nonsynonymous substitutions in this group. The relative increase in ω in hystricognaths ranged from approximately 33% in *ELAPOR1* to 95% in *Ins* (Fig. 2).

**Fig. 1.**
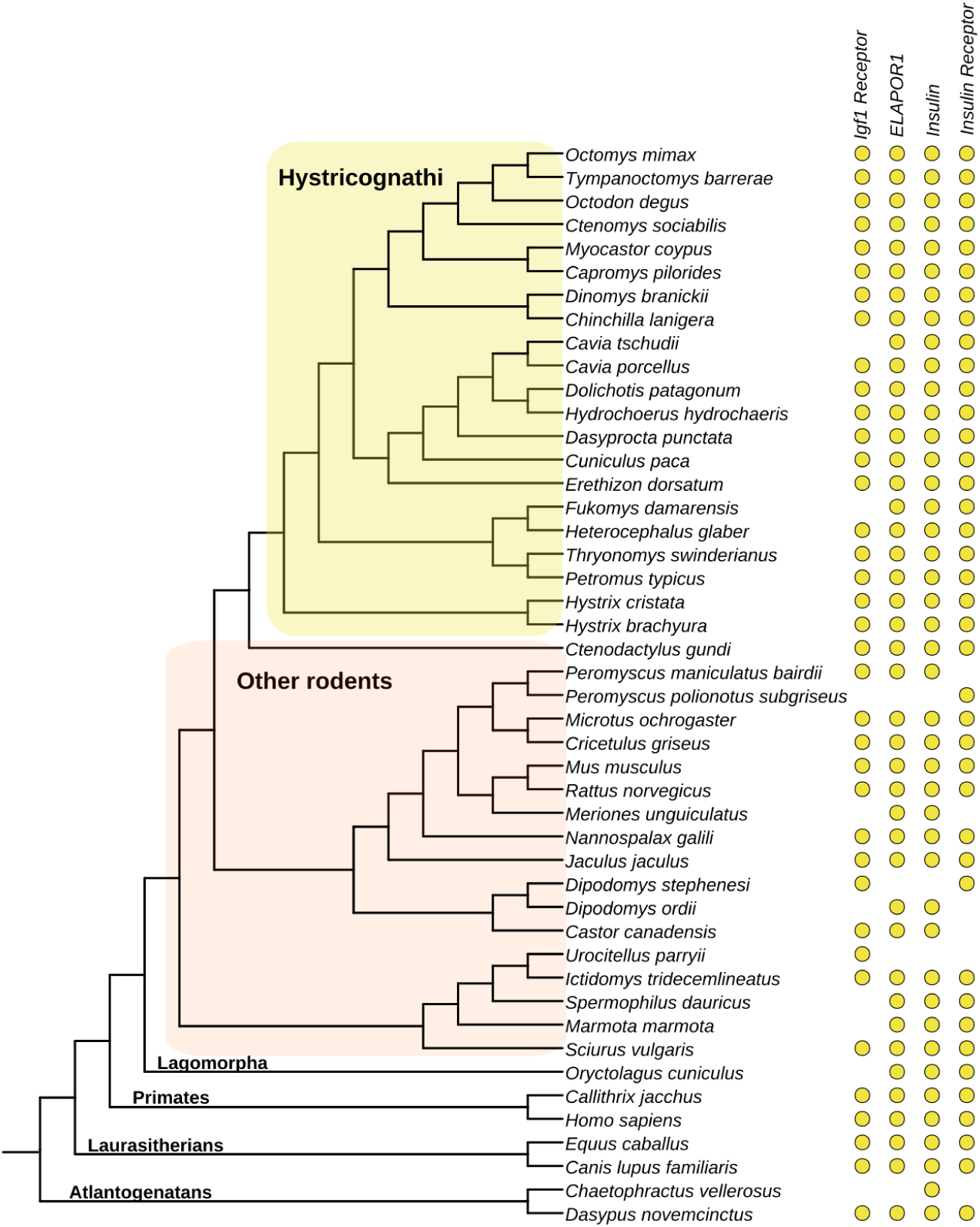
Phylogenetic relationships of the species included in this study. Tree topology is based on published literature (D’Elía et al. 2019; Upham et al. 2019). *Igf1 Receptor: Insulin-like growth factor 1 receptor; ELAPOR1: Endosome-Lysosome Associated Apoptosis and Autophagy Regulator 1*. The dots indicate the species representation for each of the loci.

**Fig. 2.**
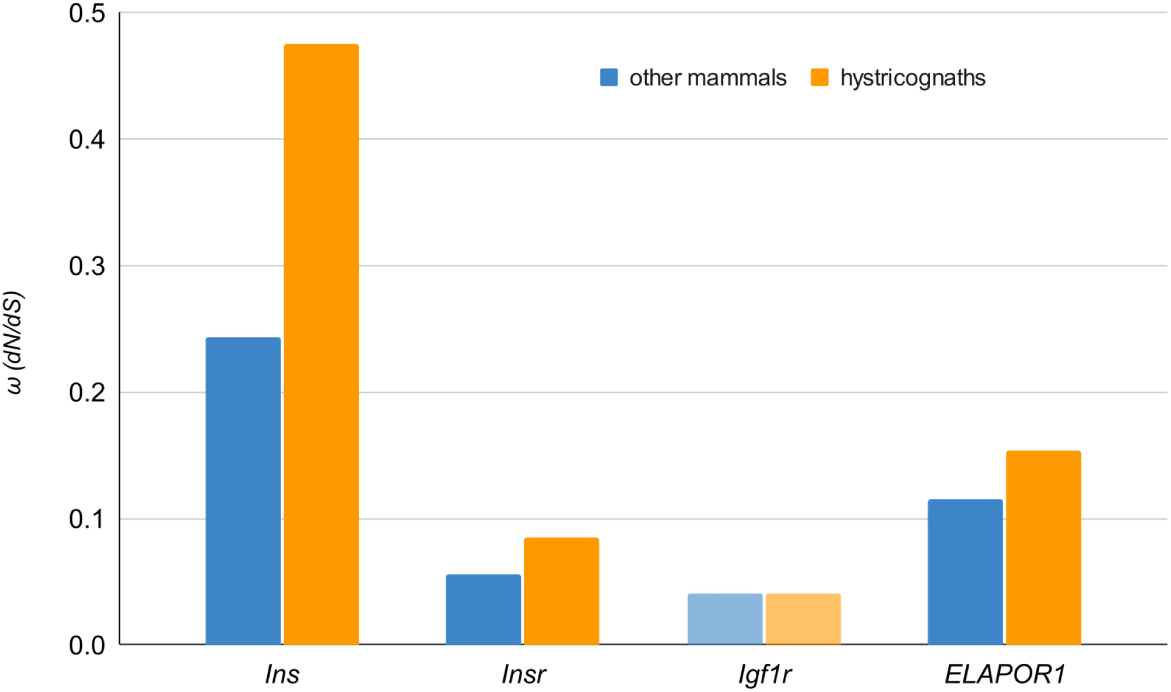
Estimates of ω for each gene based on branch models allowing two distinct ω values: one for hystricognath rodents (as a total group) and another for all other mammals. For *Igf1r*, the two-ω model did not provide a significantly better fit than the null model (*p* > 0.97). In contrast, for *Ins*, *Insr*, and *ELAPOR1*, models estimating two ω values fit the data significantly better than those estimating a single ω across all taxa (*p* < 0.00001; see Table 1).

**Table 1.**
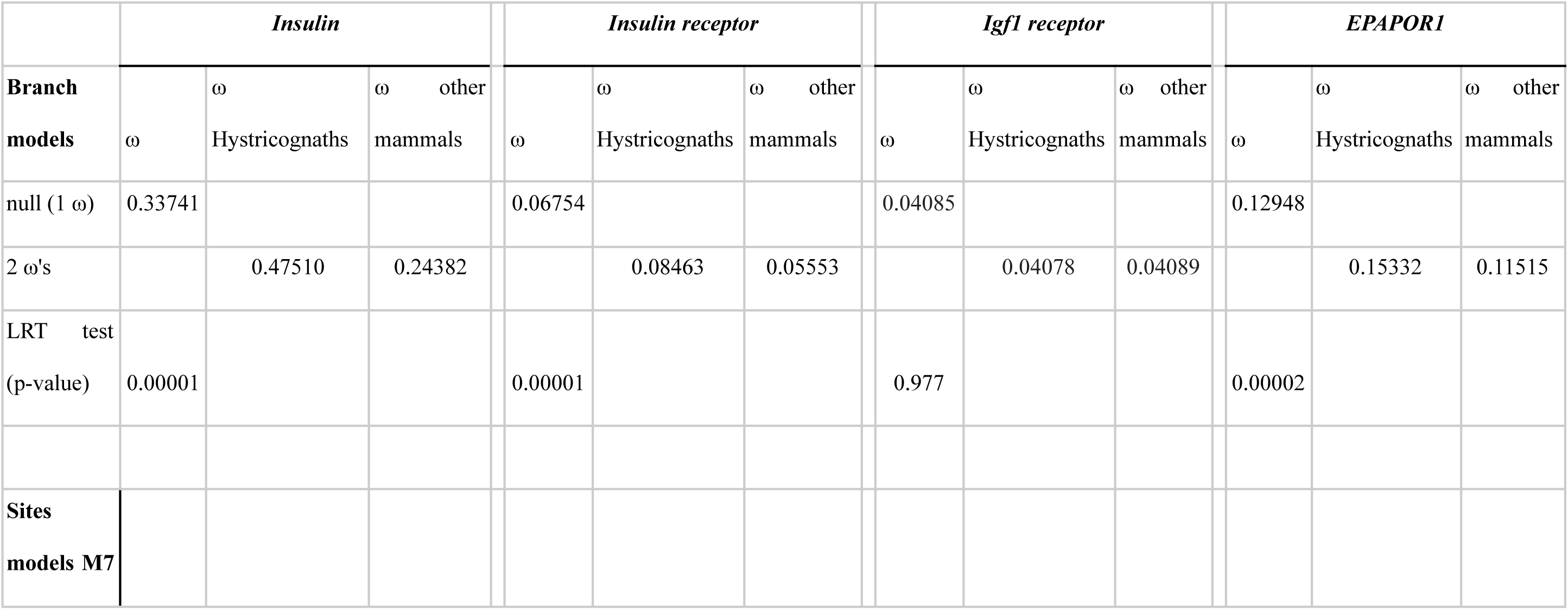
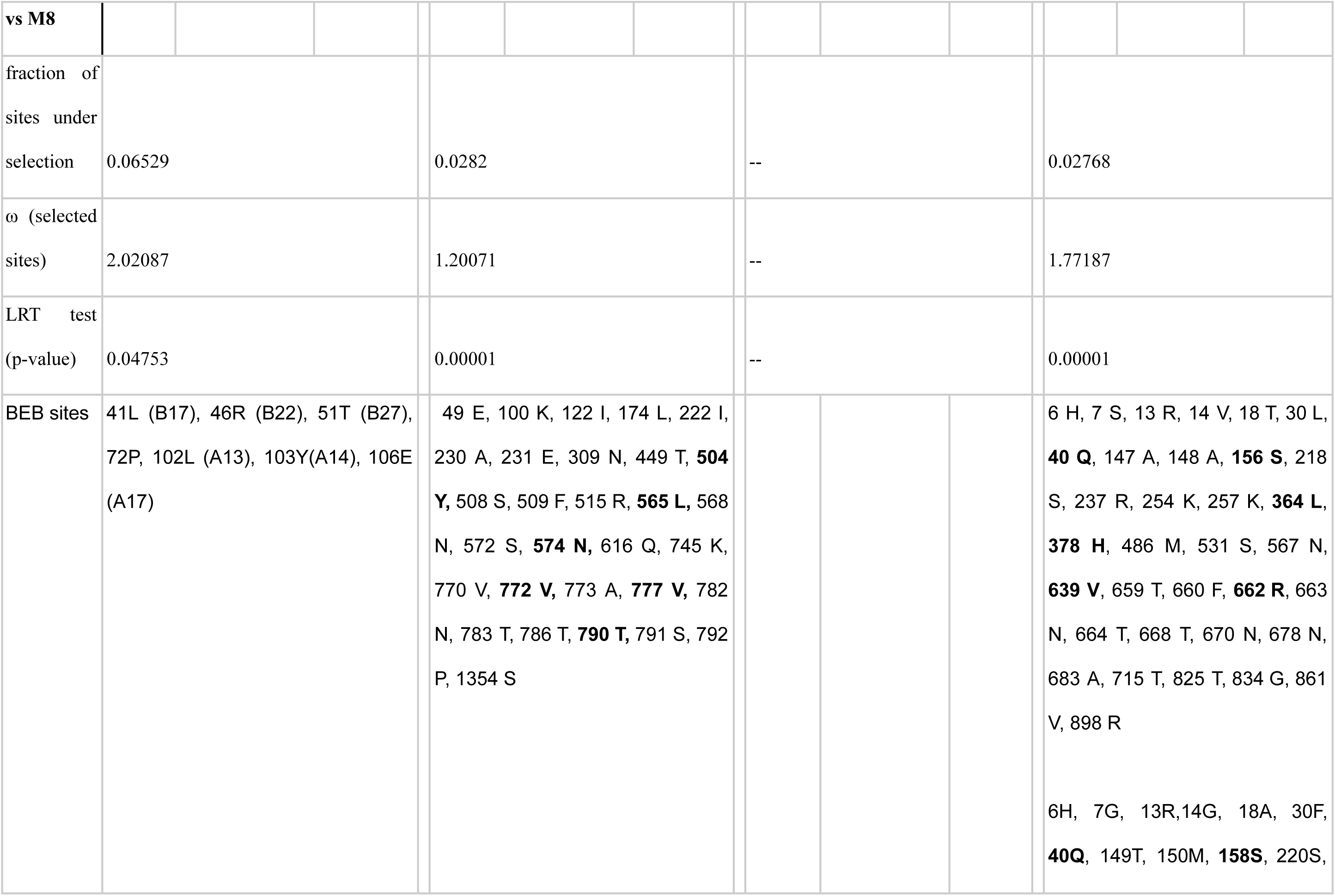
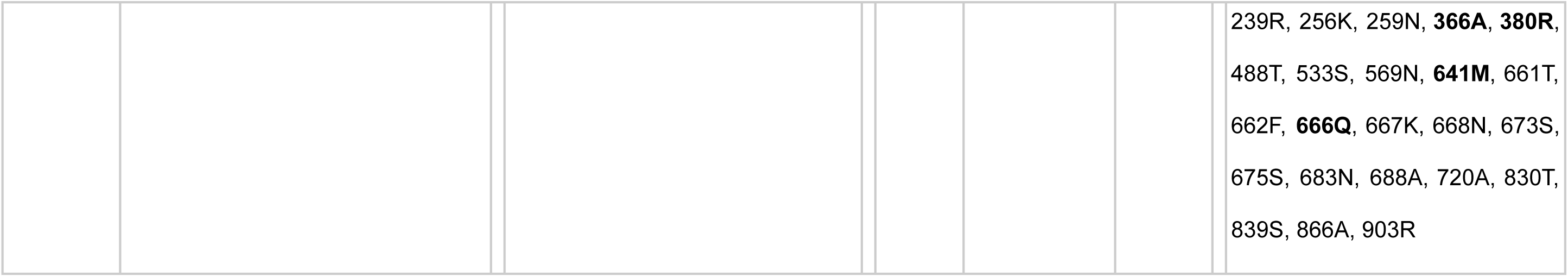
Summary of branch and site model comparisons and parameter estimates used to test for positive selection in four genes of the insulin signaling pathway. Branch models compared a null model assuming a single ω value across all mammals with an alternative model estimating two ω values: one for crown hystricognath rodents and another for all other mammals. Site models were applied only to genes that showed significant differences in ω between hystricognaths and other mammals under the branch models — that is, all genes except Igf1r. Sites with posterior probabilities p>0.95 are shown in bold. Sites are numbered according to human reference sequences (Ins: P01308; Insr: P06213; Inceptor/ELAPOR1: Q6UXG2-1). For Ins, sites located in the A and B chains of the mature insulin protein are indicated in parentheses).

Site models M7 and M8 were used to investigate further the elevated ω values observed in *Ins*, *Insr*, and *ELAPOR1*. These models differ in that M8 includes a class of sites with ω > 1, allowing for the detection of positive selection, whereas M7 does not. A significantly better fit of M8 over M7 provides statistical support for positive selection acting on a subset of codons. These analyses were conducted on datasets restricted to hystricognath rodents, focusing only on the genes that previously showed significant differences between the null branch model and the two-rate model (i.e., *Ins*, *Insr*, and *ELAPOR1*). Consequently, site models were not applied to *Igf1r*, as no significant difference was detected in the branch model comparison for this gene.

For all three genes, M8 models provided a significantly better fit than M7 models, with *p* < 0.00001 for *Insr* and *ELAPOR1*, and *p* < 0.0475 for *Ins*. The Bayes Empirical Bayes (BEB) analysis revealed variation in both the estimated ω values and the proportion of codon sites inferred to be under positive selection. Specifically, 2.8% of sites were identified as positively selected in both *Insr* (ω = 1.201) and *ELAPOR1* (ω = 1.772), whereas 6.5% of sites in *Ins* were assigned to this category, with a higher ω estimate (ω = 2.021).

### Location of sites under positive selection in the protein structures

Seven sites in preproinsulin were identified as belonging to a class under positive selection (Table 1). Of these, three sites (B17, B22, and B27) are located on the B chain, one site (codon position 72) on the C-peptide, and three sites (A13, A14, and A17) on the A chain. Notably, sites B17, A13, A14, and A17 are part of the second receptor-binding surface of mature insulin (Fig. 3; De Meyts 2015; Gutmann et al. 2020; Choi and Bai 2023). In addition, site B17 plays a critical role in hexamer formation, a key mechanism for insulin storage, together with sites B22 and A13 (Weiss and Lawrence 2018). Site B27, while not directly implicated in hexamer assembly in other mammals, is located adjacent to hexamer-forming regions in the insulin three-dimensional structure. Furthermore, adjacent sites have been modified by insertions and deletions in the families Echimyidae, Octodontidae and Ctenomyidae (Opazo et al. 2005).

**Fig. 3.**
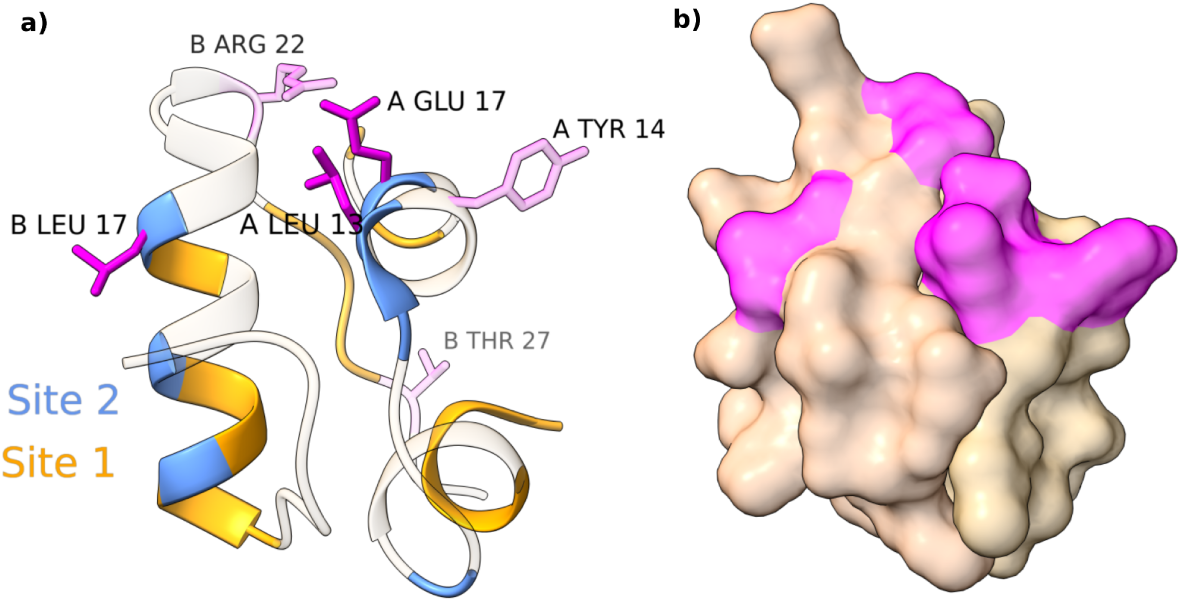
Locations of *Ins* codon sites identified as being under positive selection in hystricognath rodents. a) Sites under selection (in magenta) mapped onto the three-dimensional structure of mature human insulin. The letters preceding each amino acid (from the reference sequence PDB6o17 · INS_HUMAN) indicate whether the residue belongs to the A or B chain. b) Volume representation, with sites under selection in magenta.

Sites under positive selection in *Insr* are predominantly concentrated in the ectodomain, particularly within the first fibronectin type III domain (FIII-1) and, most notably, in the insert domain of the β-chain, located at the N-terminus of the second fibronectin type III domain (FIII-2b) (Fig. 4). In contrast, regions associated with the first insulin-binding site—specifically the C-terminus of the α-chain (αCT segment of the insert domain IDa) and the first leucine-rich repeat domain (L1)—harbor few or no positively selected sites. Additionally, no positively selected sites were identified in the third fibronectin type III domain (FIII-3). For *ELAPOR1*, positively selected sites are concentrated in the signal peptide and at the junction between the second cysteine-rich domain (C2) and the mannose 6-phosphate receptor domain (M6P) (Fig. 5).

**Fig. 4.**
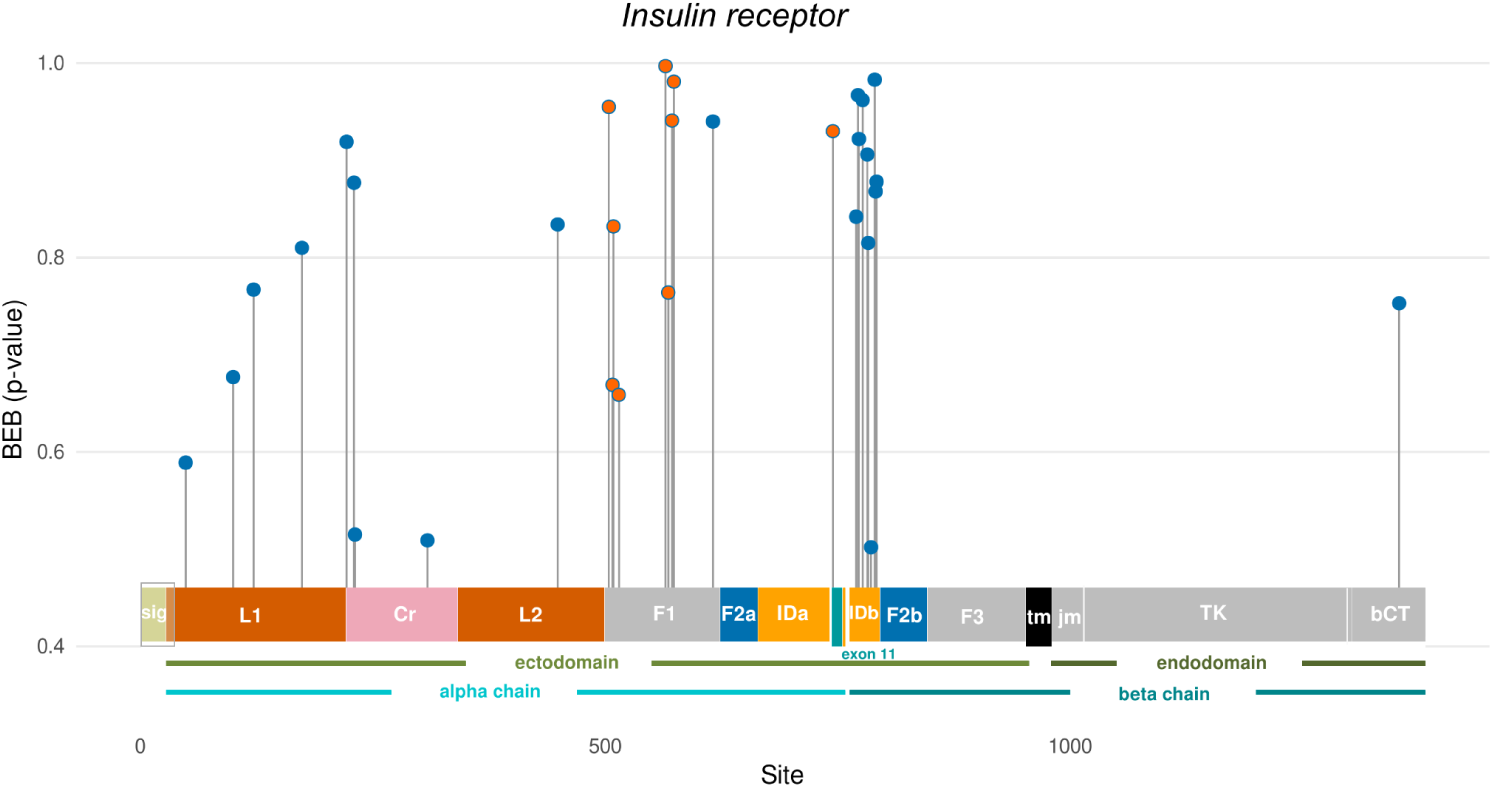
Distribution of sites under selection across the codons of *Insulin receptor*. The following regions are highlighted: sig: signal peptide; L1 and L2: first and second leucine-rich repeat domains; CR: cysteine-rich region; F1, F2 (F2a, F2b), and F3: fibronectin type III-1, III-2, and III-3 regions; IDa, IDb: insert domain, split into the alpha and beta chains; Exon11; tm: transmembrane region; jm: juxtamembrane region; TK: tyrosine kinase; bCT: beta-C-terminal (modified from UNIPROT P06213 Insr - Human). The rectangle including the cleavage peptide (27 sites) and the first 7 sites of L1 marks a region we did not include in our study because of the low quality of several sequences. Sites under selection in the second binding receptor site in F1 and in Exon 11 are highlighted in orange.

**Fig. 5.**
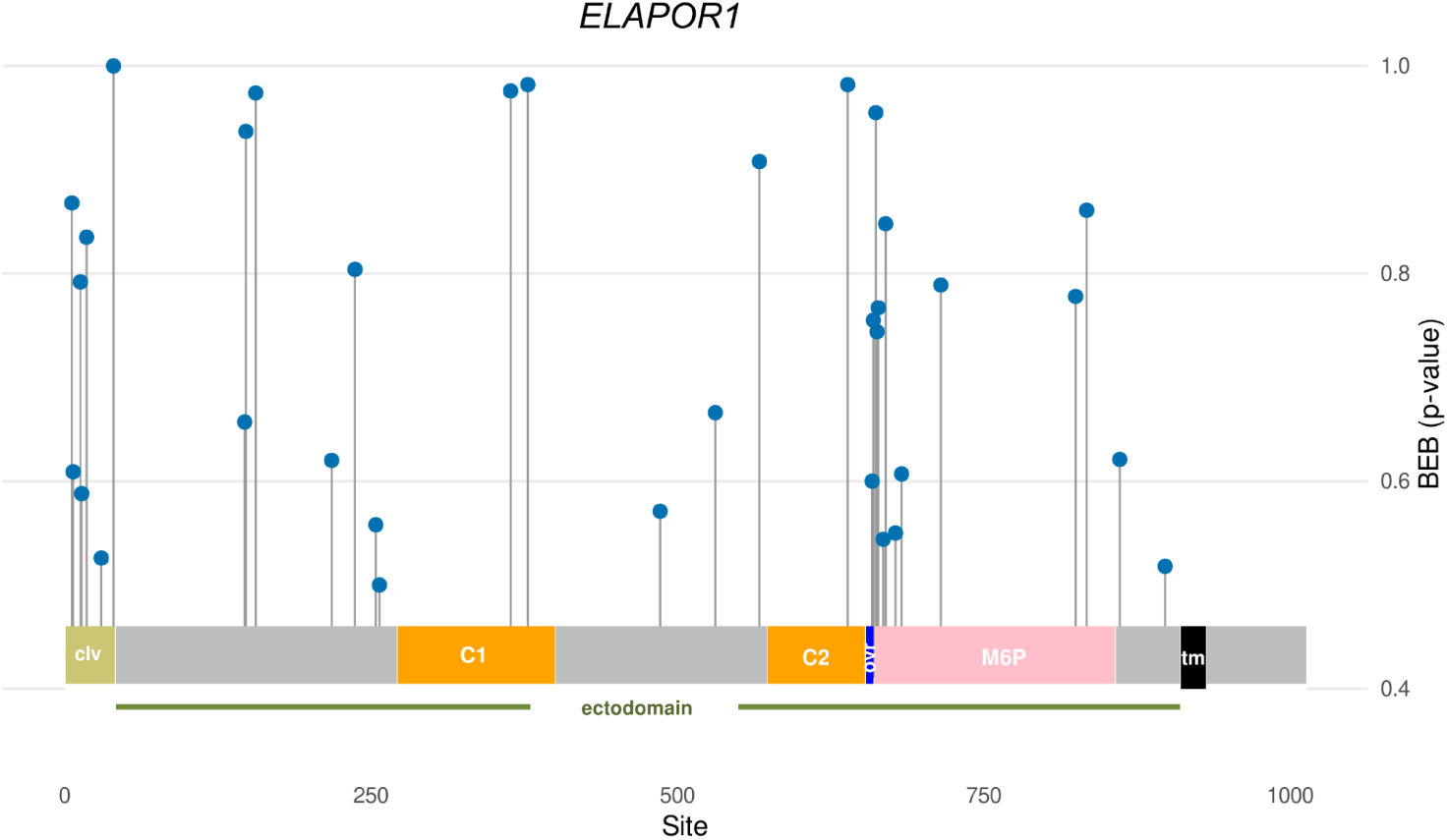
Distribution of sites under selection across the codons of *ELAPOR1*, with sites under selection in magenta. clv: cleavage (signal) peptide; C1-C2: first and second cysteine-rich domains; M6P: mannose 6-phosphate receptor; ovr: overlap between C2 and M6P; tm: transmembrane domain.

Fig. 6 compares the number of codon sites identified as evolving under positive selection (BEB sites) within specific regions of *Insr* and *ELAPOR1* to the number expected under a random distribution, based on the relative lengths of those regions. In *Insr*, the FIII-1 and FIII-2b regions contain over threefold and fivefold more BEB sites, respectively, than would be expected by chance. For *ELAPOR1*, the signal peptide harbors five times more BEB sites than expected, while the short region spanning Cys-2 and M6P displays more than 20 times the expected number, highlighting a strong signal of positive selection in this domain. In contrast, regions such as F2a, F3, the transmembrane, and endodomain regions of *Insr* are essentially devoid of positively selected sites (Figs. 4 and 6). Additional details of the results of statistical analyses are available in Supplementary Table 2.

**Fig. 6.**
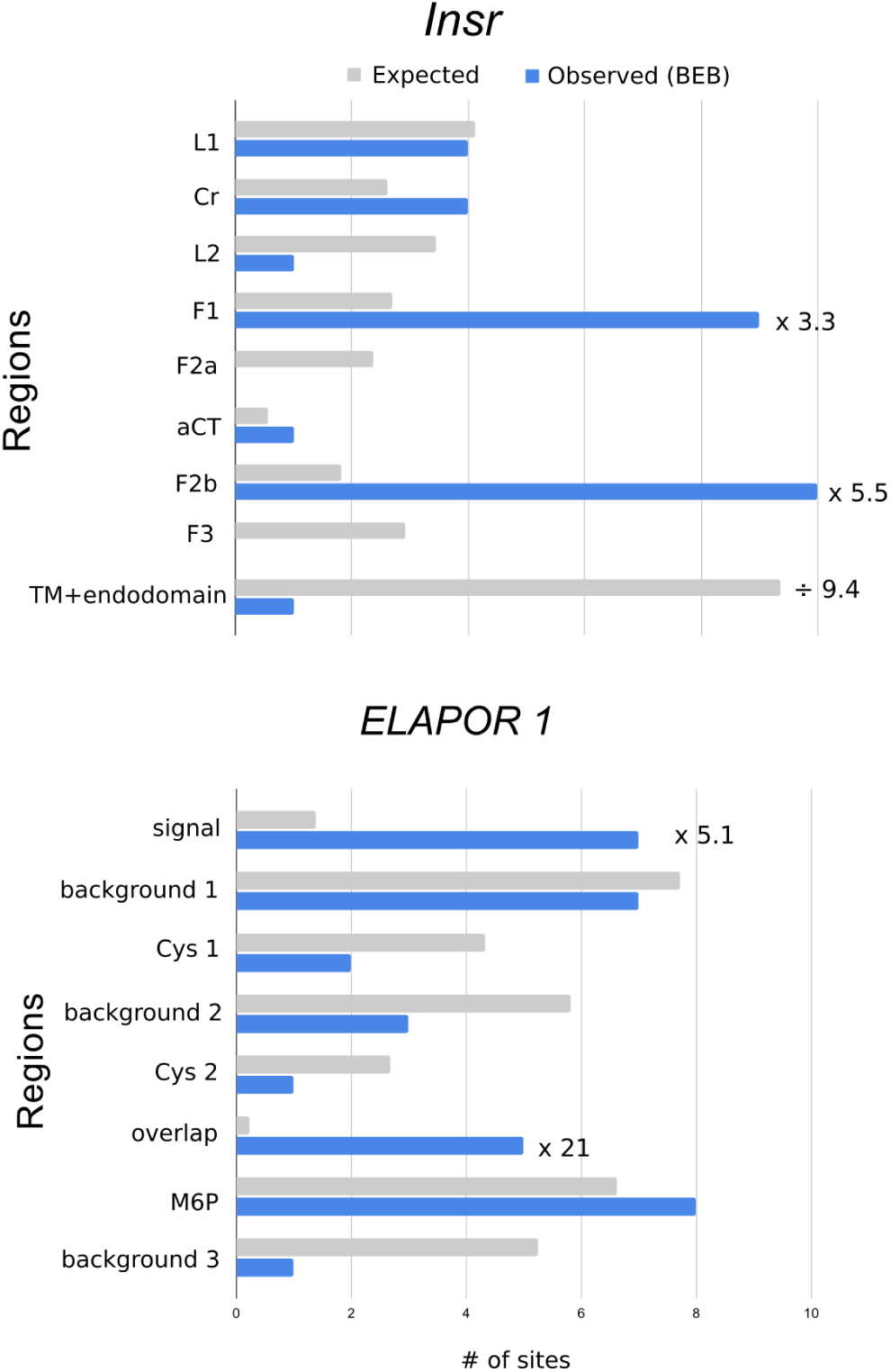
Observed number of coding sites inferred to have evolved under selection in regions of *Insulin receptor* (upper panel: L1 and L2: first and second leucine-rich repeat domains; Cr: cysteine-rich region; F1, F2 (F2a, F2b), and F3: fibronectin type III-1, III-2, and III-3 regions: aCT: alpha C-terminal; TM: transmembrane domain) and *ELAPOR1* (lower panel: Cys 1 and Cys 2: first and second cysteine-rich domains; M6P: mannose 6-phosphate receptor; overlap: overlap between C2 and M6P), compared with the corresponding expected number of sites if they were randomly distributed according to the length of each region. The ratio of observed:expected is shown for selected regions.

## Discussion

Hystricomorph rodents offer a compelling natural experiment within the Tree of Life, as several physiological systems that are typically conserved across mammals exhibit notable divergence in this group. The most striking example is the glycemic regulatory axis. In hystricognath rodents, insulin differs markedly from that of other mammals in both physiological function and amino acid sequence, and previous studies have demonstrated a role for positive selection in driving this divergence (Opazo et al. 2005). Insulin is known to bind both Insulin receptor homodimers and Insulin receptor–Igf1 receptor heterodimers (reviewed by Lawrence 2021; Choi and Bai 2023), raising the possibility that these receptors may also have evolved under selective pressure in hystricognaths. Indeed, Irwin (2020) found that ω values for *Insr* are elevated in hystricognaths compared to other rodents, although he attributed this to relaxed purifying selection rather than to positive selection. More recently, Inceptor, a newly identified regulatory protein, has been shown to modulate the insulin pathway by promoting clathrin-mediated endocytosis of Insr (and possibly Igf1r; Ansarullah et al. 2021), as well as facilitating the lysosomal degradation of cytoplasmic insulin and proinsulin within pancreatic β-cells (Siehler et al. 2024). Given the tightly integrated functional roles of Ins, Insr, Igf1r, and Inceptor, we hypothesized that the corresponding genes may have undergone positive selection in hystricognath rodents. Our analyses support this hypothesis for *Ins*, *Insr,* and *ELAPOR1* (the gene encoding for Inceptor), but not for *Igf1r*. In the following sections, we present and discuss the evidence of positive selection for each of the three genes individually.

### Insulin

In the study of Opazo et al. (2005), they included 3 of the 4 superfamilies of South American hystricognaths (the caviomorph superfamilies Cavioidea, Chinchilloidea, and Octodontoidea). Our analyses are based on a much broader representation of the diversity of hystricognaths, included all main lineages of the group. In our study we added the superfamily Erethizontoidea (New World porcupines), and several families of Old World hystricognaths, namely the Thryonomyidae (cane rats), Hystricidae (Old World porcupines), Heterocephalidae (naked mole-rat), and Bathyergidae (African mole-rats). We also significantly increased the representation of non-hystricognath rodents, including the gundi (*Ctenodactylus gundi)*, a lineage sister to the hystricognaths within the Suborder Hystricomorpha, and additional rodent lineages (Fig. 1). We also incorporated the signal and C-peptides of insulin into the analyses.

Our analyses confirm the role of positive selection in the evolution of the *Ins* gene in hystricognaths, aligning with the findings of Opazo et al. (2005). Specifically, the sites identified as being under positive selection correspond to the second binding site and/or the region involved in the formation of hexamers, as observed in other mammals. Opazo et al. (2005) identified 5 sites under positive selection, of which 3 (A13, B17 and B22) were highly significant. Those 3 sites are also recovered in our analysis with broader taxonomic representation, extending to additional families to include Old World hystricognaths. We did not recover sites B18 and B20; instead, sites A14, A17 and B27 were recovered in our analysis. Several factors may account for these differences. First, obtaining a signal of increased ω using branch models is simpler than unequivocally identifying particular sites under selection (Bielawski and Yang 2005: 120). Second, there is variation in ω across lineages; in particular, Opazo et al. (2005) showed variation among South American caviomorphs, so our inclusion of Old World taxa may include lineages with more limited adaptive divergence of *Ins*.

From a functional perspective, our findings are largely consistent with those of Opazo et al. (2005), as the positively selected sites are primarily located in the second insulin binding site and in regions associated with hexamer formation in other mammals. The only site under selection not related to those two functions in our results is B27, which is related to the first binding site (Gutmann et al. 2020) and adjacent to B26, which is involved in hexamerization in other mammals (Weiss and Lawrence 2018). There are documented insertion/deletion events in the evolution of this region in some hystricognath rodents (Opazo et al. 2005).

We also report site 72 of preproinsulin, located on the C-peptide, as being under selection. The physiological roles of the C-peptide are less known than those of insulin but include responses of endothelial tissues and the kidney (e.g., Yosten et al. 2014; Yosten and Kolar 2015; Rossiter et al. 2022). This site is spatially separated from the others in the three-dimensional structure of preproinsulin, and its functional significance has yet to be determined.

### Insulin receptor

Irwin (2020) documented increased values of ω in *Insr* of hystricognath rodents and explored the role of changes in levels of purifying selection to account for that observation. Our analysis of the evolution of this receptor shows strong evidence of adaptive evolution of this receptor. This was possible because we were able to annotate the insulin receptor gene in a large number of genomes. In addition, there has been remarkable progress in the structural analysis of the functional, dimeric insulin receptor (reviewed by De Meyts 2015; Lawrence 2021; Choi and Bai 2023). The mammalian insulin receptor is expressed as two main isoforms: Insr-A and Insr-B, which differ by the inclusion of exon 11 in Insr-B, adding 12 additional amino acids. In both isoforms, the primary transcript undergoes proteolytic cleavage to generate alpha and beta chains, which are ligated by multiple disulfide bonds. The mature receptor is a dimer composed of two such monomeric units.

The classical receptor binding site 1 of *Insr* was resolved approximately a decade ago (Menting et al. 2013, reviewed by De Meyts 2015) and involves a surface formed by the αCT and L1 regions of the alpha chain. In our analysis, we identified Lys718, a residue encoded at the start of exon 11, as having evolved under positive selection in hystricognaths. This site, located at the C-terminus of the alpha chain, interacts with insulin residue B25 of the mature insulin, which is part of its first binding site.

The most notable concentration of sites under positive selection in *Insr* occurs in two distinct regions. First, eight of the nine selected sites in the FnIII-1 domain are part of receptor binding site 2, as characterized by Gutmann et al. (2020), and appear as two clusters in the linear structure (Fig. 4): a) human residues Tyr477, Ser481, Phe482, and Arg488 located on a beta sheet of FnIII-1; and b) human residues Leu538, Asn541, Ser545, and Asn547 found on interstrand loops of the same domain. In the dimeric receptor structure reported by Gutmann et al. (2020), these sites are associated with the insulins binding to the receptor site 2 (Fig. 7). This is a significant finding, as it shows that natural selection has shaped the sequences of both the second binding site of insulin and the corresponding region in the receptor, highlighting their evolutionary interplay.

**Fig. 7.**
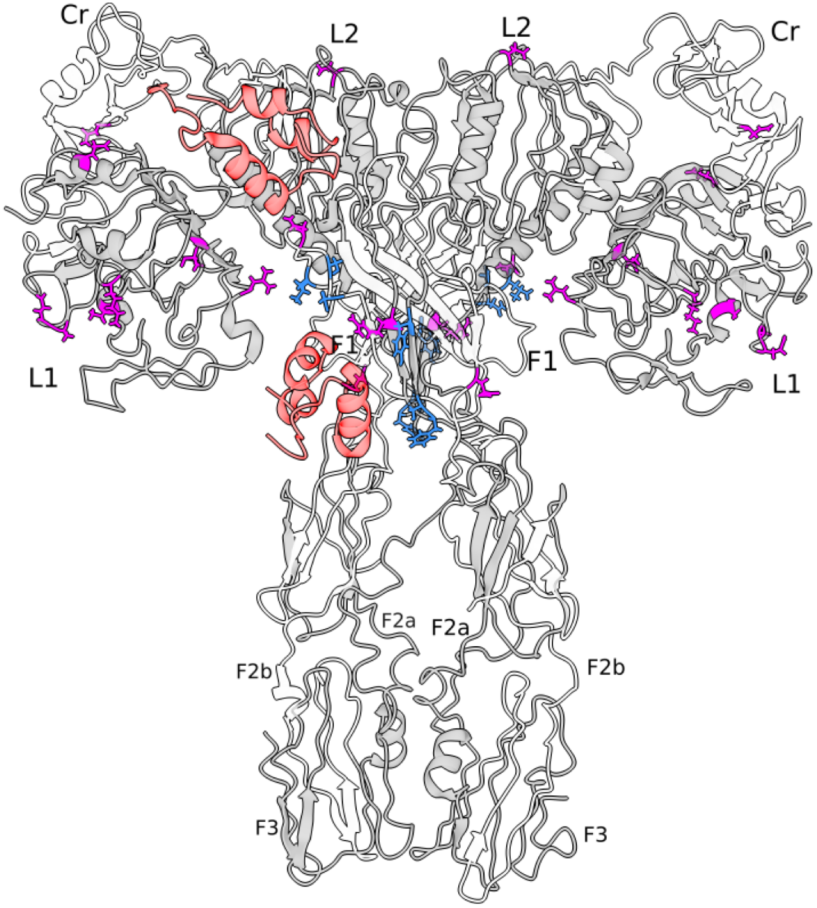
Three-dimensional structure of the mature, dimeric ectodomain of the insulin receptor, as reconstructed by Gutmann et al. (2020). Two of the four insulins that bound to the receptor (one for each type of binding site) are represented in orange. Sites under selection in the second binding domain are in blue, and the remaining ones are in magenta. This structure does not include the insert domain of the beta chain (see Fig. 5 for sites under selection in that region). L1 and L2: first and second leucine-rich repeat domains; Cr: cysteine rich region; FIII-1, FIII-2, and FIII-3: three fibronectin type III regions.

Second, ten additional selected sites are located on the βCT region, corresponding to the N-terminus of the beta chain (Fig. 4). This region, part of FnIII-2, resides along the vertical part of the active “T-shaped” structure of the ectodomain. However, due to the conformational instability of this region in the ectodomains used for structural analyses (Gutmann et al. 2020), its functional significance remains unresolved at present.

### Inceptor

Inceptor was shown to interact with the insulin receptor (Insr) and insulin-like growth factor 1 receptor (Igf1r) in the cell membrane, promoting clathrin-mediated endocytosis of Insr, thus reducing beta cell response to secreted insulin (Ansarullah et al. 2021). More recently, Siehler et al. (2024) showed that Inceptor also directs insulin and proinsulin within the beta cells towards lysosomal degradation. Thus, Inceptor appears to be a major player in the regulation of insulin in pancreatic n cells (see commentaries by Burchfield and James 2021; Hall and Choi 2021; Kulkarni 2021; and Starling 2021).

To our knowledge, this is the first study demonstrating the role of positive selection in the evolution of *ELAPOR1.* We currently lack structural analyses of the interactions of Inceptor with insulin, proinsulin, and the insulin receptor, but we identified a concentration of sites under positive selection within the signal peptide and the boundaries between the second cysteine-rich domain and the mannose-6-phosphate receptor region (Figs 5 and 6). This finding suggests changes in the regulation of *ELAPOR1* in hystricognath rodents. Hystricognaths are known not to store insulin in its hexameric form (reviewed by Weiss and Lawrence 2018) and instead secrete large quantities of insulin directly into the bloodstream (Zimmerman et al. 1974; Bajaj et al. 1986). Such physiological differences may be accompanied by changes in regulatory proteins like Inceptor. However, experimental data about levels of expression of inceptor and its interactions with proinsulin, insulin, and the insulin receptor are needed to meaningfully compare these observations with those from mouse and human models (Ansarullah et al. 2021; Siehler et al. 2024).

In summary, our analyses support the adaptive divergence of the insulin gene of hystricognaths and demonstrate that adaptive changes in the second binding site of insulin are mirrored by adaptive changes in the corresponding region of the insulin receptor. Inceptor, a recently discovered and functionally important regulator of the insulin pathway that interacts with both insulin and its receptor also shows signatures of positive selection, particularly in its signal peptide and the boundary of its second cysteine-rich region and the mannose-6-phosphate receptor. In contrast, we found no evidence of positive selection acting on Igf1r, despite its known ability to form heterodimers with Insr and to interact with Inceptor. These findings highlight that much remains to be understood regarding the evolutionary divergence of the insulin signaling pathway in hystricognath rodents.

## Materials and methods

### DNA sequences

We obtained the complete protein-coding sequences of insulin (*Ins)*, insulin receptor (*Insr)*, insulin growth factor 1 receptor *(Igf1r)*, and insulin receptor regulator (*Inceptor*, encoded by *ELAPOR1)* from the Ensembl v. 107 (Dyer et al. 2025) and National Center for Biotechnology Information (NCBI) public databases (Sayers et al. 2025). We supplemented automatic gene predictions with manual annotations of genomes using the whole-genome shotgun contigs (WGS) database from NCBI. Our sampling encompassed representative species from all major lineages of hystricomorph rodents, including bathyergids, heterocephalids, hystricids, caviids, octodontids, chinchillids, and eretizontids. Additionally, we included the sister group to hystricomorphs (ctenodactylids), as well as non-hystricomorph rodents (supramyomorphs and sciuromorphs) and non-rodent mammals, such as lagomorphs, primates, laurasiatherians, and atlantogenatans (Supplementary Table S1). For each locus, alignments were obtained using the software MUSCLE v. 5 (Edgar 2022) and inspected visually to check for consistency in interpreting intron-exon boundaries and indels. The taxa and tree used in this study are presented in Fig. 1; sequence alignments are available in Supplementary data 1.

### Natural selection analysis

We used CODEML (Yang 2007) to test for positive selection in each gene across hystricognath rodents. Specifically, we applied branch models to estimate ω values (the ratio of nonsynonymous to synonymous substitutions) under two alternative scenarios. In the null model, a single ω was estimated for all branches in the phylogeny. In contrast, the alternative model estimated two distinct ω values: one for hystricognath rodents (as a total group) and another for all remaining branches. Model fits were compared using likelihood ratio tests (LRTs). To ensure convergence, each model was run three times with different initial ω values (0.5, 1, and 2).

In addition, we used site models to examine the role of natural selection in shaping the evolution of our genes of interest. Specifically, we implemented models M7 and M8 in CODEML (Yang 2007) and compared them using LRTs. A significant LRT supported the inclusion of a class of sites with ω > 1, which is permitted in M8 but not in M7. When a significant LRT was observed, we applied Bayes Empirical Bayes (BEB) analysis to identify specific codon sites potentially evolving under positive selection. As with the branch models, each site model was run three times with different initial ω values (0.5, 1, and 2) to check for convergence.

For each locus, codons identified as being under positive selection were mapped onto the primary structure of the reference transcript to assess their distribution across functional domains. To visualize their spatial context, we used ChimeraX v1.9 (Meng et al. 2019) to map these sites onto the following three-dimensional protein structures: preproinsulin (P01308 · INS_HUMAN), insulin (PDB6O17 · INS_HUMAN), insulin receptor preprotein (P06213 · INSR_HUMAN), the insulin receptor ectodomain (EMDB: EMD-10273), and ELAPOR1 (ELAPOR1; Q6UXG2).

## Supplementary material

Supplementary Table S1: sequence and genome accession numbers.

Supplementary Table S2: results of statistical analyses.

Supplementary Data 1: sequence alignments

## Acknowledgments

This work was partially supported by a Postdoctoral Fellowship from the Agencia Nacional de Investigación e Innovación (ANII, Uruguay) to Melisa Magallanes and matching funds from the Universidad de la República (Uruguay). Juan C. Opazo acknowledges the support of the Fondo Nacional de Desarrollo Científico y Tecnológico of Chile (FONDECYT 1250688).

